# The identification of 110 novel *Mycobacterium* species within 57,974 UK Health Security Agency clinical samples

**DOI:** 10.64898/2026.11.19.767610

**Authors:** Robert J Massey, Matthew Colpus, Hieu Thai, Jeremy Westhead, Robert D Turner, Philip Bejon, Esther Robinson, Dona Foster, Colin Brown, Richard Myers, Zamin Iqbal, Philip W Fowler, Derrick W Crook, Timothy EA Peto

## Abstract

Non-tuberculous mycobacteria (NTMs) are a large diverse group of opportunistic pathogens. The accurate identification of species within this genus is vital as pathogenicity, epidemiology, and treatment differ substantially between taxa. Whole-genome sequencing has improved taxonomic resolution, however current reference databases are likely incomplete.

We analysed 53,170 high-quality mycobacterial genomes from 57,974 UK Health Security Agency samples collected between 2010 and 2025, a collection more than four times larger than the 12,726 mycobacterial genomes represented in GTDB release r232. High-quality isolates were assigned by average nucleotide identity (ANI) to recognised species within the Genome Taxonomy Database (GTDB) using a 95% ANI threshold. Isolates that fell below this threshold were clustered using a minimal dominating set approach to identify genomically coherent groups consistent with being candidate novel species. Phylogenetic analysis of all *Mycobacterium* species was used to place the observed UK diversity in a global context and assess the ecological structure within the genus.

Of the 850 Mycobacterial species-level reference genomes in GTDB, 204 GTDB references received at least one provisional assignment. We identified 586 (1.1 %) high-quality isolates that could not be assigned to any GTDB species. Genomic clustering identified 110 additional candidate species-level references, expanding the reference set from 850 to 960 (12.9%). These candidate species were broadly distributed across the phylogeny. Within the *Mycobacterium* genus we found strong evidence that phylogenetic structure is associated with human (Fritz and Purvis D = 0.10, p<0.001) and environmental sources (D = 0.34, p<0.001). Together, existing and novel reference genomes resolved the UKHSA collection into 275 species-level clusters, of which 110 (40.0%) were not represented by existing GTDB references. Furthermore, source associations differed substantially be-tween phylogenetic clades (*χ*^2^ = 364 for human and *χ*^2^ = 258 for environmental association; both p<0.001). Rarefaction analysis of the 275 species observed in UKHSA clinical samples did not plateau, with Chao1 estimation suggesting that additional sampling may reveal further novel species (Chao1 = 461).

This study provides a comprehensive analysis of genomic NTM diversity within a large national surveillance collection. Our findings suggest that NTM diversity remains incompletely characterised, and supports the value of continued surveillance and curation to improve bacterial reference databases.

## Introduction

Non-tuberculosis *mycobacteria* (NTMs) are a highly diverse group of predominantly environmental organisms, with an increasing number of NTM species recognised as opportunistic human pathogens ^1,2^. The accurate identification of NTM organisms within a clinical context is important as species differ in pathogenic potential, epidemiology, and recommended treatment ^1,3^.

Whilst whole-genome sequencing has greatly improved the consistency and resolution of bacterial classification ^4,5^, our understanding of NTM diversity remains incomplete. New species continue to be described, and existing species often have limited numbers of samples ^6,7^. Sampling efforts are concentrated on clinical cases and thus environmental diversity is likely far greater than currently observed. Consequently, our reference collections are unlikely to represent the breadth of NTM diversity that could potentially be encountered in routine clinical surveillance.

The Genome Taxonomy Database (GTDB) is a standardised genome-based framework for bacterial and archaea taxonomy based on average nucleotide identity (ANI) and alignment fraction (AF) ^8^. For defining species clusters, GTDB uses a 50% AF and 95% ANI threshold, though the ANI threshold may be higher for some species. The most recent release includes 4,967 NTM and 7,759 *M. tuberculosis* (TB) genomes collected globally from environmental and clinical sources, and defines 850 species ^8^ (of which only 6 had an ANI threshold above 95% with the highest being 95.5%). However, many species are poorly represented with 531 species (62.5 %) only being represented by a single isolate and 567 only having a placeholder name. This paucity of data constrains our understanding of within species variation and suggests that additional species are yet to be identified.

Large-scale clinical sequencing programmes, such as those undertaken by the UK Health Security Agency (UKHSA), provide an opportunity to assess how effectively existing reference databases capture the diversity encountered in routine clinical practice. These collections contain genomically coherent clusters that extend beyond recognised species boundaries. Combined with phylogenetic analysis and environmental metadata, this enables candidate species to be placed within an evolutionary and environmental context. Finally, the scale of these collections allows estimates of sampling completeness to assess whether species discovery is near saturation.

Here, we analysed 57,974 high-quality Illumina-sequenced mycobacterial clinical isolates from the UKHSA, the largest collection of NTM genomes to date. We assigned isolates to species using the GTDB and then identified genomically coherent clusters of samples that fell below the accepted 95% ANI species boundary for all currently described taxa. These clusters represent candidate novel NTM species absent from GTDB and other databases. Finally, we constructed a genus-wide phylogeny and integrated publicly available metadata to place the UK collection in the context of worldwide NTM diversity and assess the relationship between phylogeny and environment within the *Mycobacterium* genus.

## Materials and methodology

### Data collection

We analysed two datasets of mycobacterial isolates, both collected by the UK Health Security Agency (UKHSA) and its predecessor organisations: 7,794 culture-positive samples submitted to the Birmingham Mycobacterial Reference Laboratory from 2010 to 2018; and 52,761 samples collected nationally from 2018 to 2025. Duplicate results for individual samples were removed, keeping the result with the latest associated date. One sample from 2006 was removed as this was 4 years outside of the range of our other samples, and thus the associated date was assumed to be an error. Additionally, only those dated and with evidence of mycobacterial reads, according to Kraken2, were retained, giving a total of 57,974 isolates (Fig. 1). Note that *Mycobacterium tuberculosis* isolates were selectively removed by the UKHSA from the latter set, meaning that all *M. tuberculosis* isolates in this study were obtained prior to 2018 as part of the collection from the Birmingham Reference Laboratory.

**Figure 1.**
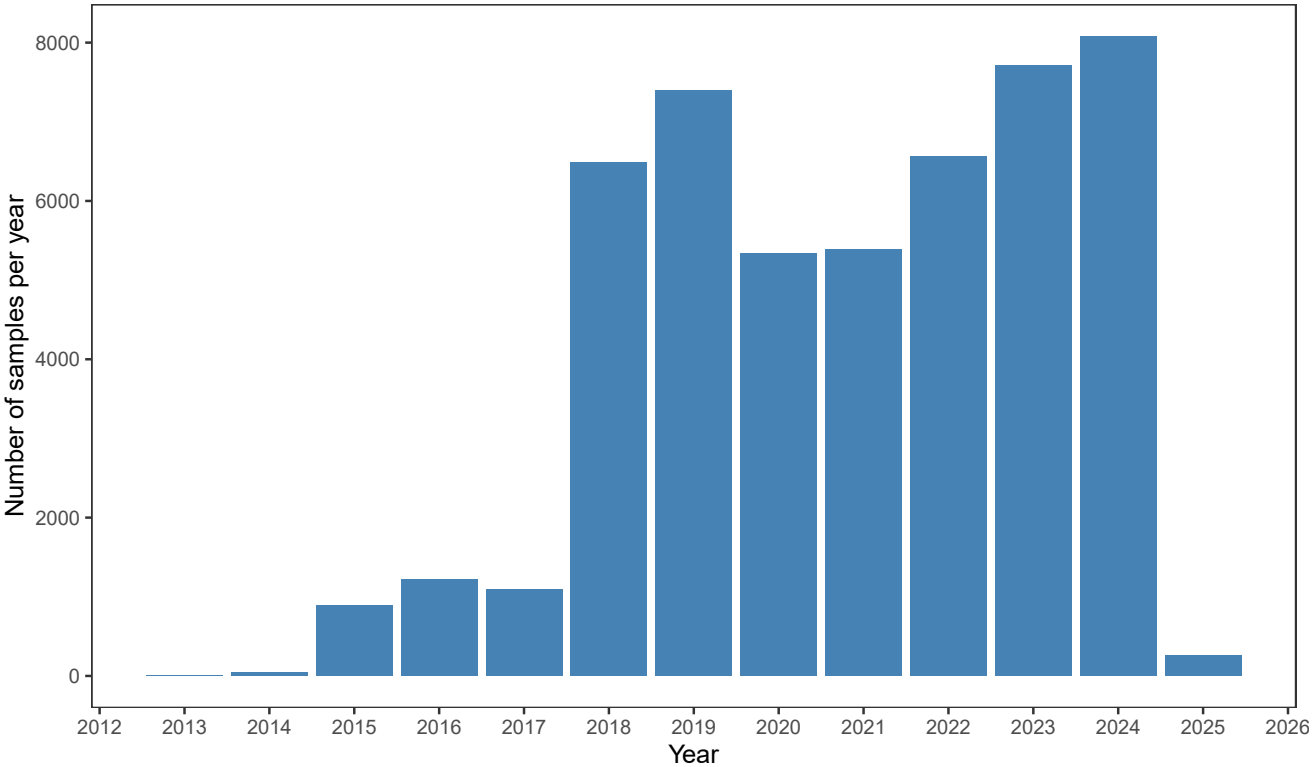
Histogram showing the number of NTM samples collected by the UKHSA between 2010 and 2025. Note that the samples collected between 2010 and 2013 were only *M. tuberculosis* samples.

### Laboratory procedures and sequencing

Libraries for Illumina sequencing were prepared using the Illumina Nextera XT protocol with manual library normalisation ^9^. Samples were batched at 12 to 16 per flow cell and sequencing was performed with the MiSeq, HiSeq or NextSeq platform.

### Genome assembly and quality control

Samples were pre-processed using a cloud-based pipeline ^10^. Human reads were removed with Hostile (v1.1.0) ^11^ or Deacon (v0.7.0) ^12^, followed by quality filtering using Fastp (v0.24.1) ^13^. Reads assigned to the *Mycobacte-riaceae* family were identified with Kraken2 (v2.1.3) ^14^ (Standard index, 6/5/2023). Reads were then assembled with Shovill (v1.1.0) ^15^ and assembly quality was assessed using CheckM2 (v1.1.0) ^16^; assemblies with *>* 80% completeness and < 10% contamination were retained for downstream analyses. Genome-based species assignment was performed using skani (v0.3.0) ^17^ to estimate ANI and alignment fraction (AF) against the GTDB ^8^ reference genomes database (r232), which includes 850 *Mycobacterium* reference genomes. In addition, GTDB-Tk ^18^ was also used to classify sample taxonomies. Finally, 221 reference type strains were previously cultured and sequenced using Illumina and were used to compare naming. The same completeness and contamination criteria were applied to these assemblies.

### Species assignment and designation of novel clusters

Skani was used to compute pairwise ANI and AF between all high-quality mycobacterial assemblies and GTDB mycobacterial reference genomes ^17^. A sample was deemed to have a ‘species-level’ assignment if it achieved *≥*95% ANI and *≥*50% alignment fraction (AF) to a GTDB reference genome. A provisional species assignment was made based on the reference with the highest ANI that met the ‘species-level’ criteria; otherwise, the reference with the highest ANI was selected. Samples were excluded if they had a provisional assignment that was not mycobacterial. Samples that did not have a skani result but were classified as belonging to the *Mycobacterium* genus by GTDB-Tk were retained but given no provisional assignment.

All samples that were not assigned a reference at the species-level were clustered in order to identify potential novel species. Pairwise ANI values were computed using skani in an all-versus-all comparison. A graph was then constructed in which nodes represented samples and edges connected pairs of samples sharing *≥*95% ANI and *≥*50% alignment fraction (AF). A minimum dominating set (MDS) was identified using the graph and defines the smallest set of reference genomes required for all isolates to be assigned at a species-level. Where multiple MDS solutions existed, we selected the set that maximised the mean ANI between isolates and their assigned references. This yielded the minimal number of additional references required for complete species-level assignment, a reference set we refer to as ‘GTDB+’. The new species clusters were provisionally designated NOVEL_NTM_1, NOVEL_NTM_2, etc. in descending order of cluster size. Having created these additional putative references, all samples were assigned a final species designation based on the reference genome with the highest ANI.

### Phylogenetic tree construction

We constructed a phylogenetic tree for the `GTDB+` reference genomes (n=960) along with six members of the *Mycobacteriales* order (*Corynebacterium diphtheriae, Gordonia bronchialis, Nocardia farcinica, Rhodococcus erythropolis, Segniliparus rotundus*, and *Tsukamurella paurometabola* (see Table S3) to serve as outgroups, as has been done previously ^19^. We used gtdbtk (v2.7.2) ^18^ to identify and align 120 bacterial marker genes. Maximum-likelihood inference was performed with IQTree (v3.1.2) ^20^, which was run on the concatenated alignment using the settings `-alrt 1000 -bb 1000 -m MFP` and the ModelFinder selected the `Q.INSECT+F+R10` substitution model. The resulting tree was rooted using the outgroups.

The tree was divided into clades using TreeCluster ^21^ (`‘max_clade’` method) based on patristic distances, using a threshold of 0.1 to separate clades. The patristic distance between two species in the tree is the sum of the branch lengths along the path connecting the two tree tips. Additionally, NCBI metadata was collated, for all *mycobacterial* samples used in GTDB, to classify isolate source and geographical location, where possible. This was used to investigate the correlation between phylogeny and ecological niche. After removing species with only one isolate, and then further removing clades with less than 5 member species, corrected odds ratios and Chi squared tests were preformed to test the association between the constructed clades and ecological niches. Similarly, phylogenetic association with ecological or geographic niche was assessed using Fritz and Purvis’ *D* statistic, with each region/niche being an independent test. Independently of the inferred phylogeny we also assessed these associations using pairwise amino-acid (AA) p-distances calculated from the concatenated GTDB 120 bacterial marker genes, excluding positions containing gaps or ambiguous residues in either genome. AA p-distance is equivalent to one minus the pairwise AA identity for this region. Only high quality genomes were used for these analyses. For human and environment source, a binary distance matrix was constructed (0 = same association status; 1 = different status) and compared against the AA p-distances using Pearson Mantel tests with 9,999 permutations. Local clustering was further assessed by grouping genome pairs into 20 BAC120 amino-acid p-distance half-open bins of width 0.01, spanning 0 to 0.20. For each bin, discordance was calculated as the proportion of pairs with different source status. For Human, Environment, Land, and Aquatic source status again a binary distance matrix was constructed (1 = same association status; 0 = different status), and source labels were then shuffled across genomes without replacement 9,999 times, preserving the observed prevalence; discordance was recalculated for each bin after every shuffle.

### Rarefaction analysis

Rarefaction analyses were conducted on all high-quality mycobacterial assemblies. To assess the completeness of NTM species diversity captured by the dataset, species accumulation curves were constructed using a permutation-based approach. For each of 100 permutations, samples were randomly ordered and cumulative species richness recorded as each successive sample was added. Mean richness and 95% quantile intervals were calculated across permutations at every sample size. Total species richness was estimated using the Chao1 non-parametric estimator, implemented via the vegan R package ^22^.

### Statistical analysis

Chi-squared tests were used to assess the heterogeneity of ecological association between the clades constructed for the phylogenetic tree. The R package `caper` ^23^ was used to calculate the Fritz and Purvis D statistic ^24^. The R package `vegan` ^22^ was used to perform the Pearson Mantel statistical tests. All statistical analyses were performed in R v4.5.1^25^. Analysis code will be made publicly available on submission.

## Results

After filtering with Kraken2, samples contained a median of 1.13 million (IQR 0.78-1.75 million) total reads. Of the initial 56,776 assembled genomes, 1,782 (3.14%) were excluded because they were incomplete (completeness *≤* 80%), and 1,816 (3.20%) were excluded because they were presumed to be mixed infections (contamination *≥* 10%). A further 8 sequences were removed due to suspected contamination or assignment to non-mycobacterial organisms (Fig. 2). This yielded 53,170 high quality *Mycobacterium* assemblies for the main analysis. They had a median of 338 contigs (IQR 253-484) and a median contig N50 of 58 kbp (IQR 41-81 kbp).

**Figure 2.**
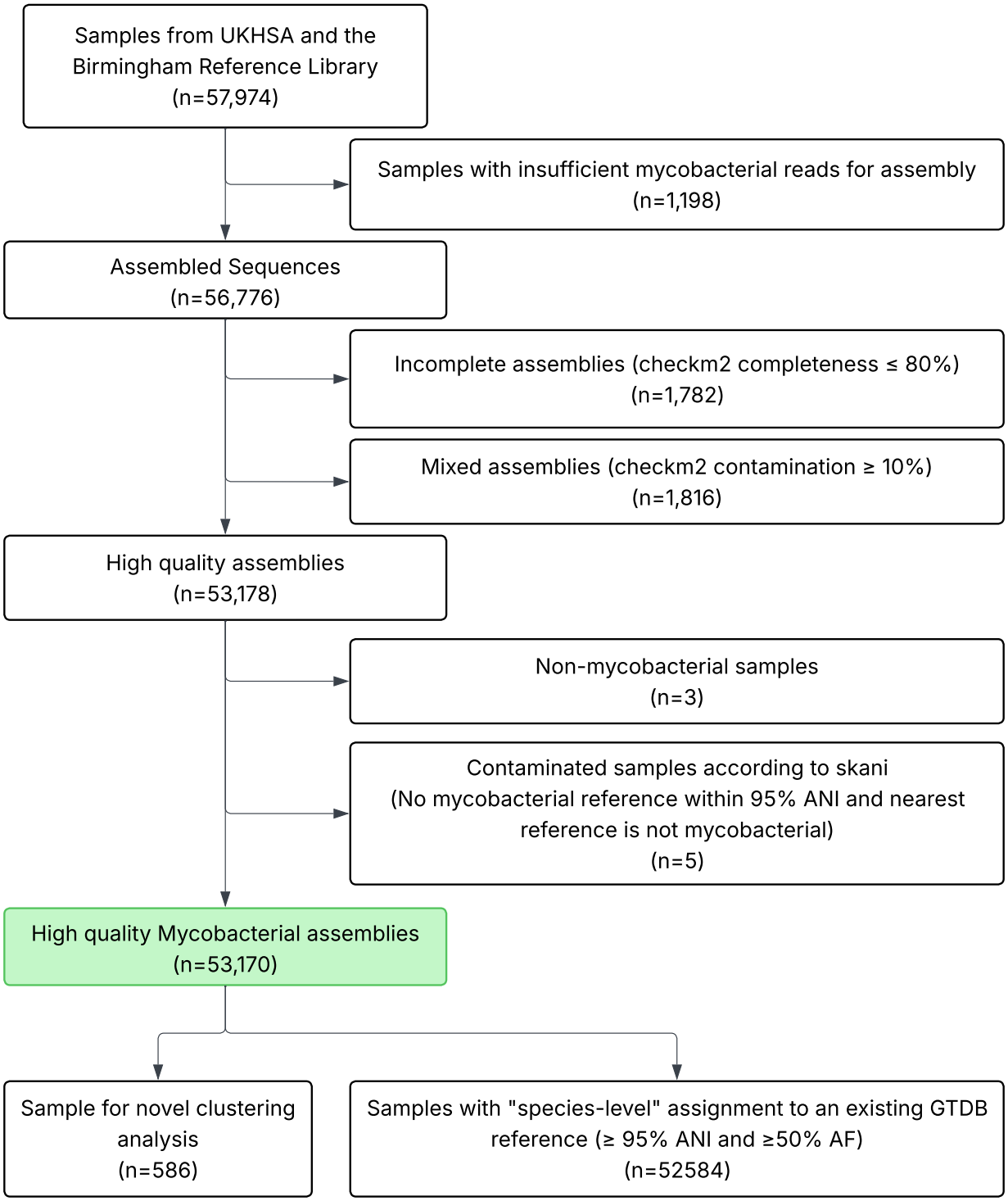
Bioinformatic filtering

### Species assignment reveals extensive novel diversity

We used ANI and alignment fraction (AF) to compare the 53,170 high-quality mycobacterial assemblies with the 850 *mycobacterial* species in GTDB (r232). Of these 850 references, 204 had at least one isolate provisionally assigned to them. Figure 3 shows the ANI distribution for the most frequently assigned, with all species shown in Fig. S1.

**Figure 3.**
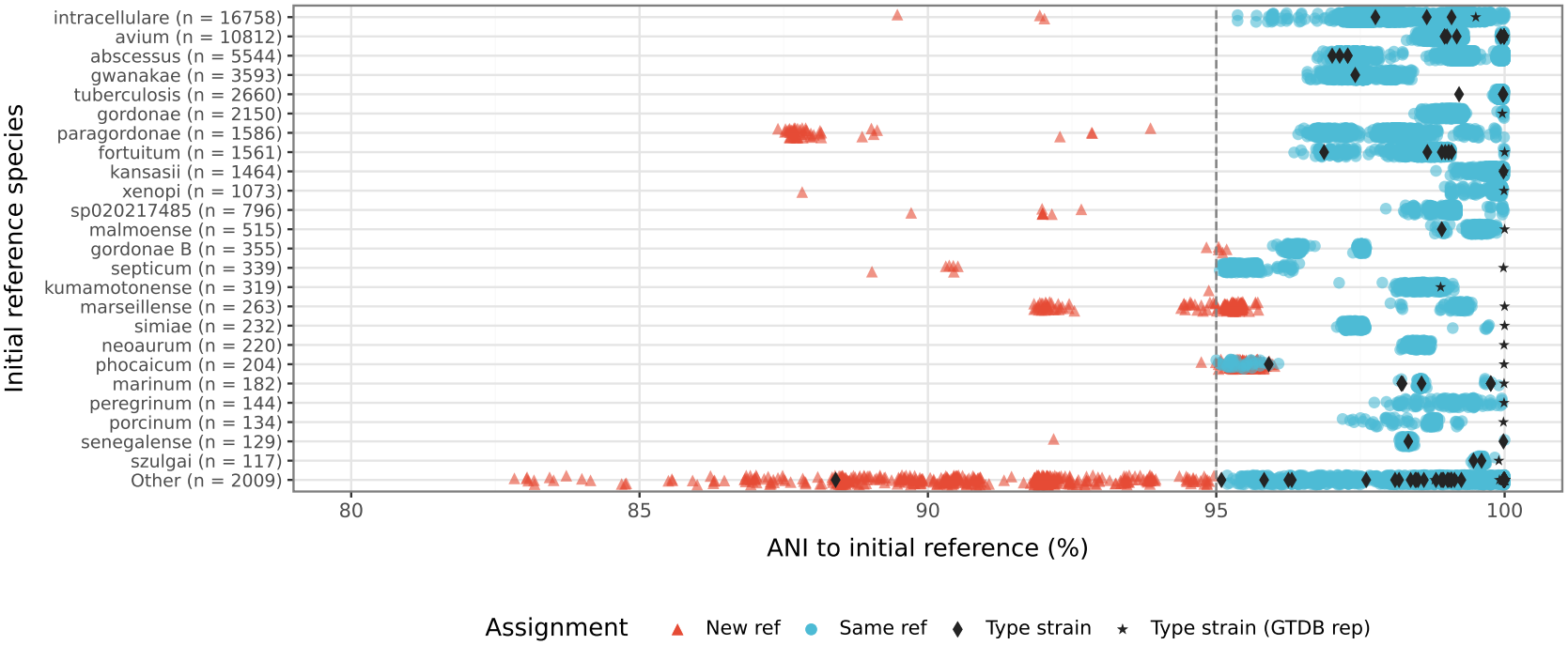
Distribution of ANI against provisionally assigned GTDB reference. The top 24 most common species are shown individually, whilst the “Other” row contains all other species. The red triangles indicate samples which were later assigned to a novel species cluster, whilst blue dots kept the provisional reference. Samples that did not achieve a provisional assignment were not included in this figure. Additionally, type strains are marked with black diamonds or black stars if they match the strain used by GTDB representative.

In total, 586 isolates did not meet the species-level assignment criteria for any GTDB reference (ANI *≥* 95% and AF *≥* 50%). We therefore used a minimum dominating set (MDS) approach to determine the minimum number of additional reference genomes required to ensure that every isolate met these criteria for at least one reference. This identified 110 isolates as new reference genomes, each representing a putative species-level taxon not represented among the 850 GTDB references. We therefore classified these lineages as 110 novel candidate species. These genomes were combined with the 850 GTDB reference genomes to form the 960-genome ‘GTDB+’ reference set. Species assignments obtained via skani and GTDB-Tk were concordant for all isolates meeting the species-level thresholds.

Reassignment against `GTDB+` yielded 275 species-level clusters, comprising 165 clusters represented by an existing GTDB reference and the 110 novel candidate species. Of the 275 clusters, 104 (37.8%) of the clusters contained a single isolate, whilst 83 (30.2%) contained at least 10 isolates. Among the 110 novel candidate species, 60 were singletons, 37 contained between 2 and 9 isolates, and 13 contained at least 10 isolates. The most common species found were *M. intracellulare* (n=16,755), *M. avium* (n=10,812), and *M. abscessus* (n=5,544). Note that *M. chimaera* is considered part of *M. intracellulare* within GTDB.

The extent to which GTDB reference genomes represented within species diversity varied among species (Fig. 3). For instance, all *M. tuberculosis* isolates were highly similar to the GTDB reference as measured by ANI. In contrast, the majority of isolates assigned to *M. septicum* and *M. phocaicum* were somewhat distant to their corresponding reference genome as indicated by having <97% ANI. Additionally, seven pairs of GTDB references had ANI *≥*95% to each other (ANI 95.00% to 95.53%, Table S1), highlighting that it is sometimes difficult to define distinct species using a universal threshold.

We additionally compared our reference type strain assemblies against the GTDB and `GTDB+` references, and used StrainInfo ^26^ to identify corresponding GTDB references where possible (Fig. 3 & S1). Two type strains, DSM 44364 (*M. ratisbonense*) and NCTC 9822 (*M. gordonae*), were assigned to the novel candidate species `NOVEL_NTM_1` (relative of *M. phocaicum*) and `NOVEL_NTM_4` (relative of *M. sp963726875*) respectively. This highlights that not all type strains have assembled genomes represented in GenBank or GTDB, and that our novel candidate species are defined relative to available reference genomes. We also identified a discrepancy for *M. kumamotonense* between our type strain (JCM 13453) and the GTDB assembly (DSM 45093), despite these records referring to the same strain.

### Human-associated mycobacteria cluster phylogenetically and genetically

We constructed a phylogeny from the 960 `GTDB+` reference genomes and used six outgroup taxa to root the tree (Methods; Fig. 4). The 110 novel candidate species were distributed across the phylogeny, with at least one in 19 of the 35 (54.3%) clades. However, this diversity was concentrated in a relatively small number of clades; for example, clades 18, 6, and 11 together contained more than half of the additional putative species.

**Figure 4.**
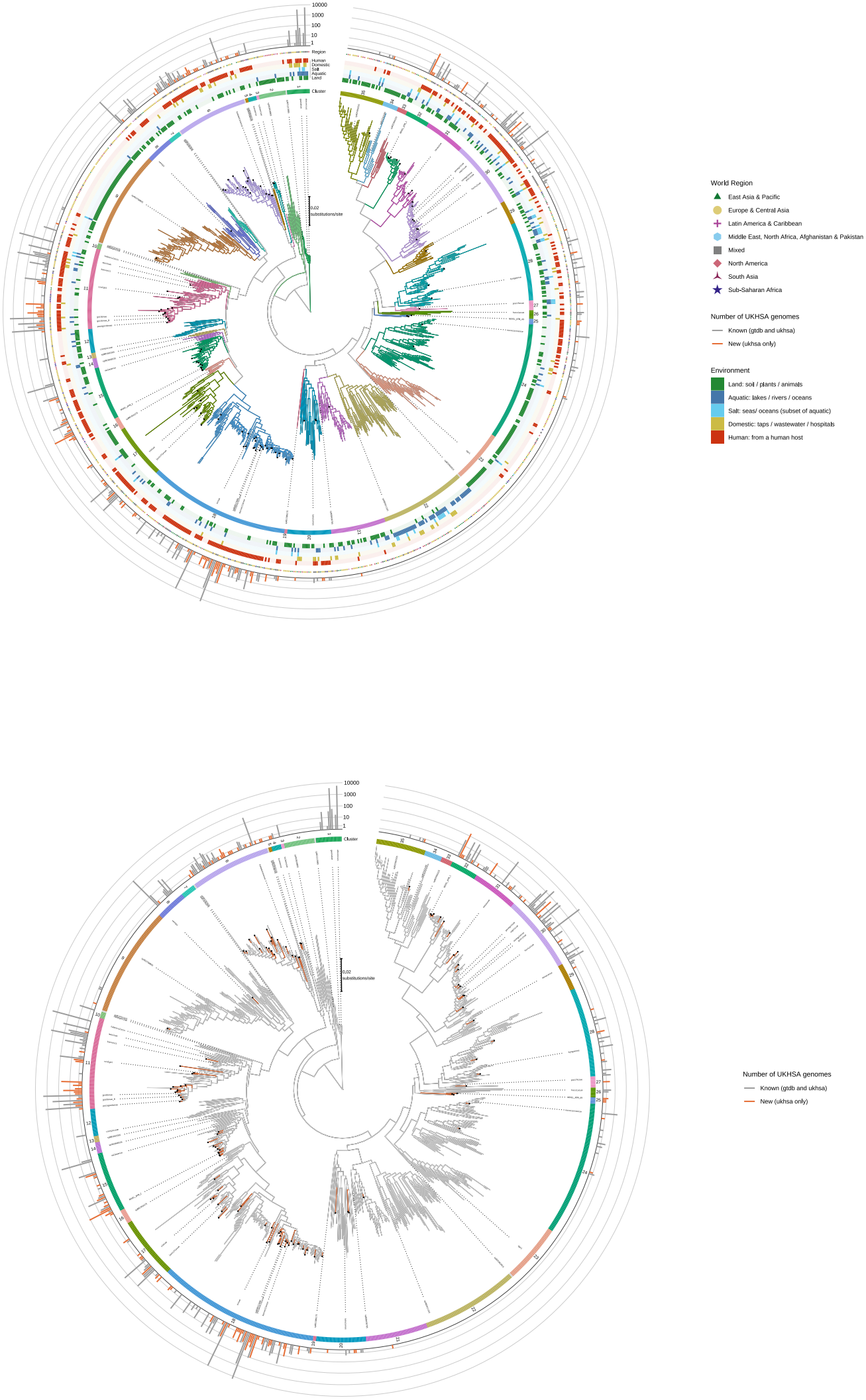
Phylogenetic tree for *Mycobacterium*. a) The branches are coloured based on clade grouping. This is also shown in the first ring. Tips with black dots indicate novel species clusters. Rings 2-4 show what environments that species has been found in. Ring 5 shows which geographic region that species has been found in. Ring 6 is a log-scaled bar chart showing the number of samples found for each species within the UKHSA collection. Novel species have an orange bar. Certain species have been labelled: those with at least 100 isolates and the largest species within each clade. b) Now branches are coloured orange for novel species. The rings for clade and UKHSA samples counts are the same as the previous panel.

**Figure 5.**
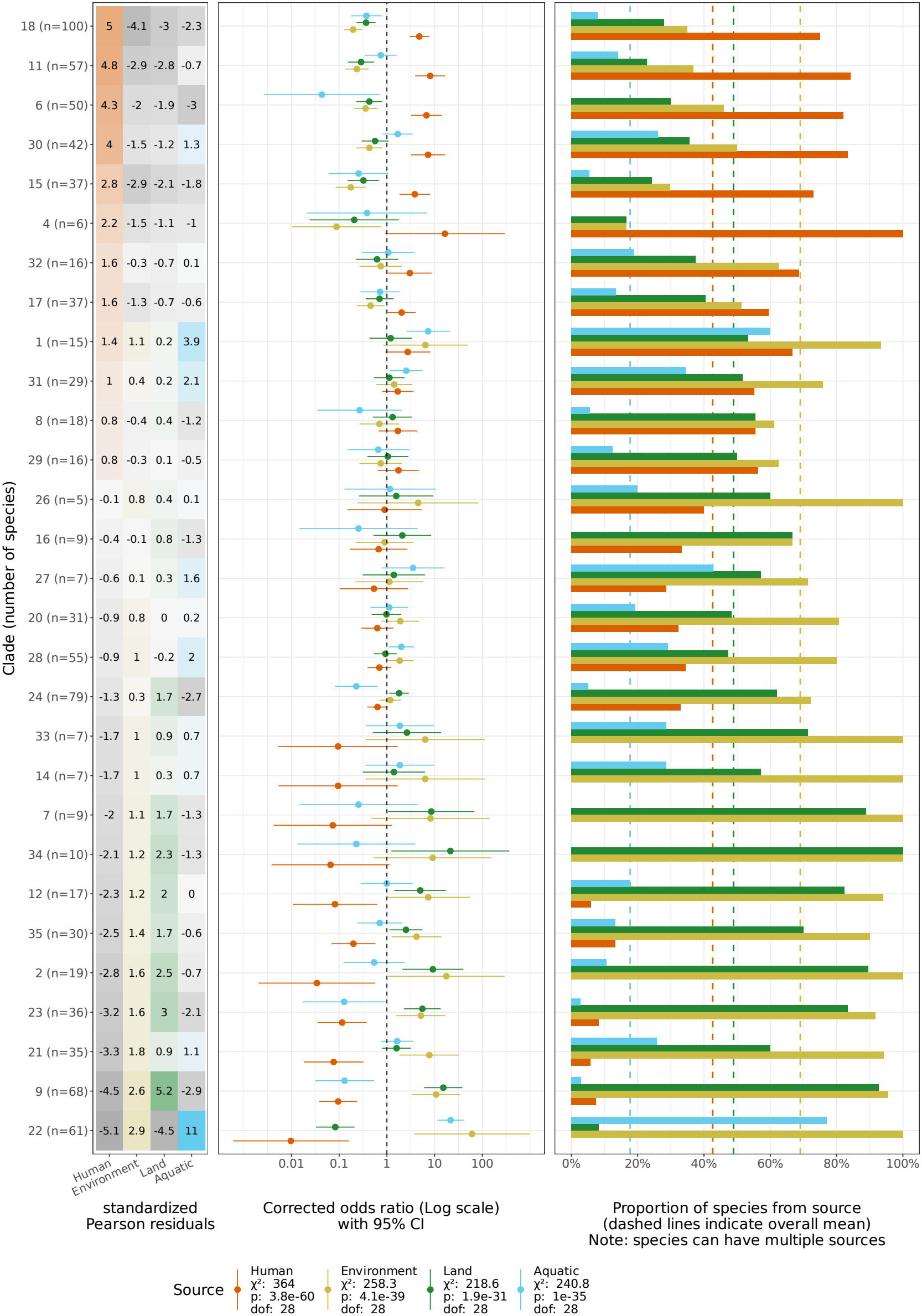
Distribution of source types within clades after removing singleton species and clades with less than 5 member species. a) Heatmap showing the standardised Pearson residuals from calculating the *χ*^2^-statistic for each source-environment. Positive values indicate greater levels of that environment than expected. Negative values indicate under-representation. b) Forest plot showing corrected odds ratio with 95% confidence intervals. Bar-chart showing the proportion of species found in each source-environment. The *χ*^2^-statistic, p-value and degrees of freedom are provided in the legend. Clades have been ordered by the residuals for “Human”. Note that a given species may be found from multiple sources. Aquatic (oceans/rivers/fish) and Land (soil/plants/animals) 12 are subsets of Environment.

We found a strong phylogenetic signal for human source association (Fritz and Purvis D = 0.10, p<0.001), consistent with the significant heterogeneity observed between clades. The phylogenetic signal for environmental source was somewhat weaker (D = 0.34, p<0.001; Figs. S4 & 5). This weaker signal likely reflects the widespread occurrence of environmental species across the phylogeny, including within clades enriched for human association. Indeed, as all NTMs are believed to be environmental, with sufficient sampling an ecological niche would be identified for all species.

Because recombination may influence the phylogenetic reconstruction, we additionally assessed source-association clustering without reference to the inferred tree. For this we looked directly at the relationship between source-association dissimilarity with the pairwise amino-acid (AA) p-distances from the concatenated marker genes (see Methods). Grouping pairwise genome comparisons into 0.01 AA p-distance bin revealed substantial local clustering: for genome pairs that had an AA p-distance between 0.01 - 0.02, 78.1% shared human-source status (50.5% expected by chance, p=0.002. Supplementary Fig. S3 and Table S5). Similarly, for environmental-source status, for those pairs in the 0.02 - 0.03 AA p-distance bin 67.4% shared environmental-association status (53.8% expected, p=0.0032). This enrichment declined with increasing genetic distance. The same relationship was observed for land and aquatic source-types. Global Pearson Mantel tests showed weak correlation for human association (r=0.056, p=0.0001) and showed no positive association for environmental status (r=-0.021, p=0.998), suggesting that the source-association clustering only applies locally. Together, these findings demonstrate that the observed phylogenetic signal is not solely an artefact of tree reconstruction.

The clade-level analyses reflected these broader patterns. Several clades were strongly enriched for human-associated species; for example, species in clade 11 had an OR of 8.1(95% CI 3.9 - 17) of being human-associated compared to species in other clades. Conversely, some clades were strongly enriched for environmental species, such as clade 22, in which species had an OR of 60.6 (95% CI 3.7 - 983) of being environment-associated (Fig. S4).

We found a much weaker phylogenetic signal for geographic region, with all regions having D*>*0.7 except south Asia (D=0.61). Interpretation was limited by biases in geographical representation: 560 species were observed in Europe and central Asia, compared to only 34 in south Asia. No clades were found to be restricted to a single region.

We next assessed how well the UKHSA collection captured the phylogenetic diversity of the *Mycobacterium* genus by measuring the evolutionary distance from each of the 850 GTDB species to its closest representative in the UKHSA collection (Fig. 6). Among human-associated species, 165/289 (57%) were represented by the UKHSA collection, and 90% were within a patristic-distance of 0.029 of a UKHSA species. In contrast, only 89/638 (14%) of environmentally associated species were found by UKHSA, and the cumulative proportion grew more slowly, reaching 90% only at a patristic distance of 0.066. These results suggest that UK clinical surveillance captures a large proportion of the diversity associated with humans, whereas substantial additional diversity would only be captured by environmental sampling.

**Figure 6.**
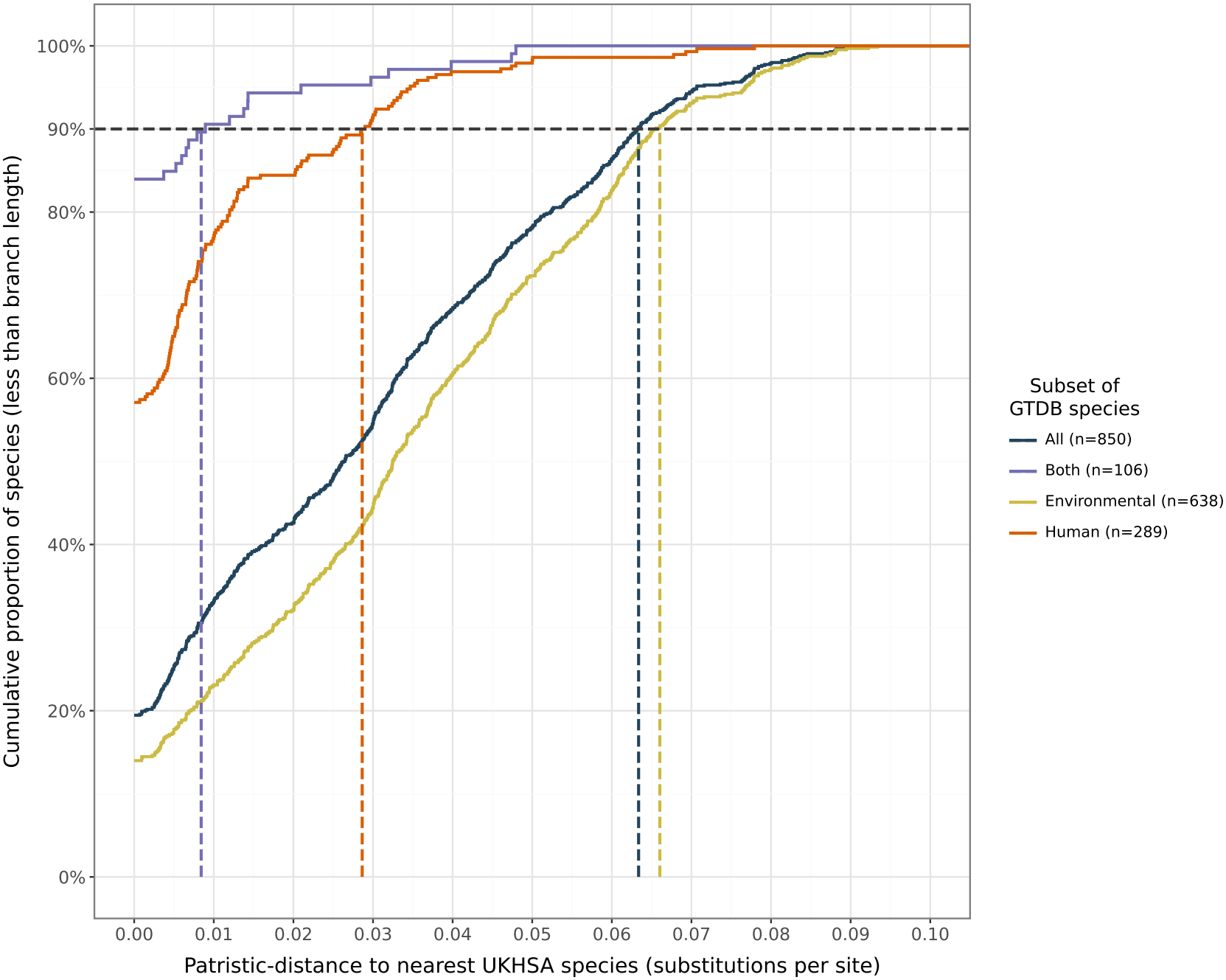
Cumulative distribution graph of distances from GTDB species to nearest species included in the UKHSA collection. Patristic distance is the sum of branch lengths between two tips (i.e. species) in the phylogenetic tree. Note that this does not include the 110 novel species identified. The value at the y-axis indicates the proportion of species whose nearest UKHSA representative is within the patristic specified distance..

### *Mycobacterium* species discovery has not reached saturation

To assess whether species discovery had approached saturation, a rarefaction curve was constructed using all high-quality *Mycobacterial* assemblies. We used our `GTDB+` species designations (n=275 observed species) to investigate how the number of species accumulates with increasing sample size, including the rate at which isolates belonging to the novel candidate species were detected.

The `GTDB+` rarefaction curve showed only slight plateauing, suggesting that additional sampling may reveal yet more *Mycobacterium* species (Fig. S5). Consistent with this, the Chao1 estimator returned an estimated total richness of 461 *Mycobacterium* species, substantially larger than the 275 species observed in the present collection. It is worth noting here that the chao1 estimate is conditional on the sampling process and taxonomic clustering scheme, and may not necessarily estimate the total number of mycobacterial species capable of appearing in a UK clinical context.

## Supporting information

Manuscript Supplementary Material

## Discussion

In this study, we have analysed the largest collection to date of *Mycobacterium* isolates, made up of 53,170 high-quality genomes (of which 2,660 were TB). Using the GTDB taxonomic framework ^8^ we resolved these genomes into 275 Mycobacterial species; while the majority of isolates could be assigned to 165 (60.0%)recognised species, we identified 110 (40.0%) genome clusters outside of the accepted species boundaries and these are therefore candidate novel species. This demonstrates that, whilst current reference databases successfully resolve the species present in most clinically relevant isolates, a substantial portion of clinical diversity remains incompletely represented.

Comparison with the complete GTDB database further highlights that the NTM isolates from the UKHSA collection are specific to a clinical context. Of the 850 GTDB *Mycobacterial* references, 685 were absent from our dataset despite its size. This suggests that UKHSA surveillance captures a subset of the overall genus diversity, representing those species within the UK capable of infecting or colonising humans. The large number of absent species likely reflects lineages rarely seen in clinical practice, confined predominantly to environmental reservoirs. Our phylogenetic analysis indicates that association with human clinical specimens is phylogenetically linked.

Several clades were highly enriched for human-associated species, whilst other clades were purely environmental. Indeed, this association was also observed directly from genetic distance, with closely related species being significantly more likely to share a source type. This pattern suggests that ecological adaptation, particularly the ability to colonise humans or persist in environments linked to human exposure, is conserved within lineages. It is worth noting, however, that our analysis did not directly assess pathogenicity. Conversely, we find geographic distribution to have much less phylogenetic signal, concordant with the global prevalence of several NTM lineages ^27^.

Although clinical diversity is phylogenetically structured, large scale surveillance studies are uniquely placed to distinguish and characterise candidate novel species, providing more evidence than is possible from a smaller dataset. Despite the size of our dataset, our rarefaction analysis suggests that species discovery within the UK clinical context is not yet at saturation. Although the most abundant 27 species (each represented by *≥*100 isolates) account for 96.3% of all samples, the species accumulation curve did not asymptote, and the Chao1 estimate predicts greater species richness than observed. Nevertheless, the majority of this additional diversity appears to be concentrated amongst rare taxa: 60 (54.5%) of the 110 novel clusters contained only one example genome, and only 13 (11.8%) contained at least ten genomes. These findings suggest that the most prevalent strains within the UK are now well characterised, and that continued surveillance is likely to recover increasingly rare or environmentally associated taxa, with diminishing returns for clinical interpretation and action.

However, continued surveillance may also provide the opportunity to refine existing species references. We identified several species, most notably *M. septicum* and *M. phocaicum*, where all samples in our collection were unusually distant (as indicated by a low ANI) from the reference. Updating reference taxa may improve the accuracy of WGS-based species identification, which in turn will provide a stronger foundation for future studies to investigate within-species variation.

Several limitations should be considered when interpreting these findings. First, our analysis was performed within the existing GTDB taxonomy and used the conventional species threshold of 95% ANI. Although this is a consistent and widely applicable framework, a universal threshold may not adequately reflect the genomic diversity of all taxa, and thus alternative thresholds may would merge or split some species clusters identified here. Indeed, we observed seven pairs of existing GTDB reference genomes with ANI *≥*95%, illustrating that species boundaries are not always consistent with a universal threshold. Furthermore, identification of additional candidate species-level clusters relied on a reference-centred classification framework, rather than direct evidence of discrete species boundaries. Genetic diversity may be continuous between species, with isolates sitting between reference genomes. This is particularly true within a complex of closely related species (e.g. the *M. avium* complex), where genomic boundaries may be difficult to resolve. As such, the additional reference genomes identified in our analyses should not necessarily be interpreted as representing biologically distinct organisms. Instead, they identify previously uncharacterised diversity that is not adequately represented in current reference sets. Phenotypic description and characterisation is therefore required before being given a formal species designation. Second, this dataset is biased as it comprises all routine clinical submissions to the UKHSA rather than systematic environmental sampling or random sampling from the population. Consequently, species frequency estimates are not directly generalisable to broader ecological settings or to clinical settings in other countries. Similarly, the rarefaction analysis presented here does not reflect true ecological species richness and should be interpreted within the context of clinical sampling. Finally, isolation of a species from a clinical sample does not necessarily imply disease and the lack of linked patient data prevents the assessment of associations between genomic variation and clinical presentation, prevalence, or patient outcomes.

In conclusion, this study demonstrates that whilst current taxonomies describe the dominant clinically relevant NTM species they still do not capture all of the genomic diversity observed within routine clinical practice. Continued large-scale genomic surveillance will therefore be important both for refining bacterial taxonomy and for maintaining bacterial databases as additional rare clinical lineages are discovered.

### Transparency Statement

PWF and DWC receive consultancy fees from Ellison Institute of Technology, Oxford Limited (EIT).

### Funding

This research is funded by the National Institute for Health and Care Research (NIHR) Health Protection Research Unit in Healthcare Associated Infections and Antimicrobial Resistance (NIHR207397), a partnership between the UK Health Security Agency (UKHSA) and the University of Oxford, the National Institute for Health and Care Research Biomedical Research Centre: Oxford (NIHR203311), the Ineos Oxford Institute for antimicrobial research and EIT. The views expressed are those of the authors and not necessarily those of the NIHR, UKHSA, the Department of Health and Social Care or EIT. For the purpose of open access, the author has applied a CC BY public copyright licence to any Author Accepted Manuscript version arising from this submission.

### Author contributions

The analysis in this manuscript was conceived by TEAP, DWC, and PWF. ER, DF, RM, CB, HT, RT, JW, MC, and RJM led and performed sequencing and bioinformatics. TEAP, MC, and RJM guided and performed the analyses in this manuscript. RJM and MC wrote the manuscript. ZI provided methodological insights that were important to the content and structure of the manuscript. All authors provided feedback on the final manuscript.

### Ethics statement

This study used mycobacterial sequences collected by UKHSA as part of routine surveillance activity. These were de-identified and shared with the University of Oxford through the NIHR Health Protection Research Unit in Healthcare Associated Infections and Antimicrobial Resistance. Public health action taken as a result of surveillance is one of the UKHSA’s key roles as stated in the Health and Social Care Act 2012 and subsequent Government directives which provide the mandate and legislative basis to undertake necessary follow-up. This work is part of service development carried out under this framework, and as such explicit ethical approval is unnecessary.

## References

[1] Falkinham, J. O. Environmental sources of nontuberculous mycobacteria. Clinics in Chest Medicine 36, 35–41 (2015).

[2] Honda, J. R., Virdi, R. & Chan, E. D. Global environmental nontuberculous mycobacteria and their contemporaneous man-made and natural niches. Frontiers in Microbiology 9, 2029 (2018).

[3] Daley, C. L., Iaccarino, J. M., Lange, C., Cambau, E., Wallace, J., Richard J. et al. Treatment of nontuberculous mycobacterial pulmonary disease: an official ATS/ERS/ESCMID/IDSA clinical practice guideline. European Respiratory Journal 56, 2000535 (2020).

[4] Quainoo, S., Coolen, J. P. M., van Hijum, S. A. F. T., Huynen, M. A., Melchers, W. J. G. et al. Whole-genome sequencing of bacterial pathogens: the future of nosocomial outbreak analysis. Clinical Microbiology Reviews 30, 1015–1063 (2017).

[5] Ferraz Helene, L. C., Klepa, M. S. & Hungria, M. New insights into the taxonomy of bacteria in the genomic era and a case study with rhizobia. International Journal of Microbiology 2022, 4623713 (2022).

[6] Gebert, M. J., Lipner, E. M., Galletta, J. M., Henley, J. B., Hoffert, M. et al. A cross-environment comparison of nontuberculous mycobacterial diversity. Applied and Environmental Microbiology 92, e01552–25 (2026).

[7] Koh, W.-J. Nontuberculous Mycobacteria – overview. Microbiology Spectrum 5 (2017).

[8] Parks, D. H., Chuvochina, M., Rinke, C., Mussig, A. J., Chaumeil, P.-A. et al. GTDB: an ongoing census of bacterial and archaeal diversity through a phylogenetically consistent, rank normalized and complete genome-based taxonomy. Nucleic acids research 50, D785–D794 (2022).

[9] Pankhurst, L. J., del Ojo Elias, C., Votintseva, A. A., Walker, T. M., Cole, K. et al. Rapid, comprehensive, and affordable mycobacterial diagnosis with whole-genome sequencing: a prospective study. The Lancet Respiratory Medicine 4, 49–58 (2016).

[10] Westhead, J., Baker, C. S., Brouard, M., Colpus, M., Constantinides, B. et al. Characterizing the performance of an antibiotic resistance prediction tool, gnomonicus, using a diverse test set of 2,663 Mycobacterium tuberculosis samples. Microbial Genomics 11, 001592 (2025).

[11] Constantinides, B., Hunt, M. & Crook, D. W. Hostile: accurate decontamination of microbial host sequences. Bioinformatics 39, btad728 (2023).

[12] Constantinides, B., Lees, J. & Crook, D. W. Deacon: fast sequence filtering and contaminant depletion. bioRxiv (2025). Doi:10.1101/2025.06.09.658732.

[13] Chen, S., Zhou, Y., Chen, Y. & Gu, J. fastp: an ultra-fast all-in-one FASTQ preprocessor. Bioinformatics 34, i884–i890 (2018).

[14] Wood, D. E., Lu, J. & Langmead, B. Improved metagenomic analysis with Kraken 2. Genome Biology 20, 257 (2019).

[15] Seemann, T. Shovill. https://github.com/tseemann/shovill (2016).

[16] Chklovski, A., Parks, D. H., Woodcroft, B. J. & Tyson, G. W. CheckM2: a rapid, scalable and accurate tool for assessing microbial genome quality using machine learning. Nature methods 20, 1203–1212 (2023).

[17] Shaw, J. & Yu, Y. W. Fast and robust metagenomic sequence comparison through sparse chaining with skani. Nature Methods 20, 1661–1665 (2023).

[18] Chaumeil, P.-A., Mussig, A. J., Hugenholtz, P. & Parks, D. H. GTDB-Tk v2: memory friendly classification with the genome taxonomy database. Bioinformatics 38, 5315–5316 (2022).

[19] Gupta, R. S., Lo, B. & Son, J. Phylogenomics and comparative genomic studies robustly support division of the genus Mycobacterium into an emended genus Mycobacterium and four novel genera. Frontiers in microbiology 9, 67 (2018).

[20] Wong, T. K., Ly-Trong, N., Ren, H., Demotte, P., Baños, H. et al. IQ-TREE 3: phylogenomic inference software using complex evolutionary models. Molecular Biology and Evolution 43, msag117 (2026).

[21] Balaban, M., Moshiri, N., Mai, U., Jia, X. & Mirarab, S. TreeCluster: Clustering biological sequences using phylogenetic trees. PloS one 14, e0221068 (2019).

[22] Oksanen, J., Simpson, G. L., Blanchet, F. G., Kindt, R., Legendre, P. et al. vegan: Community ecology package (2022). R package version 2. 6-4.

[23] Orme, D., Freckleton, R., Thomas, G., Petzoldt, T., Fritz, S. et al. caper: Comparative analyses of phylogenetics and evolution in R (2025). R package version 1.0.4.

[24] Fritz, S. A. & Purvis, A. Selectivity in mammalian extinction risk and threat types: a new measure of phylogenetic signal strength in binary traits. Conservation Biology 24, 1042–1051 (2010).

[25] R Core Team. R: A Language and Environment for Statistical Computing. R Foundation for Statistical Computing, Vienna, Austria (2025).

[26] Lissin, A., Schober, I., Witte, J. F., Lüken, H., Podstawka, A. et al. StrainInfo—the central database for linked microbial strain identifiers. Database 2025, baaf059 (2025).

[27] Hoefsloot, W., van Ingen, J., Andrejak, C., Ä ngeby, K., Bauriaud, R. et al. The geographic diversity of nontuberculous mycobacteria isolated from pulmonary samples: An NTM-NET collaborative study. European Respiratory Journal 42, 1604–1613 (2013).

