## Supplementary material for "The identification of 110 novel *Mycobacterium* species within 57,974 UK Health Security Agency clinical samples": Manuscript Supplementary Material

### 1 List of extra files

1. All samples: Table of all isolates used in the study. Provides read and assembly statistics as well as the provisional and final species assignments.
2. Rooted tree: Phylogenetic tree newick file
3. Sample source classification: Table of source/region metadata for all public samples
4. Reference source summary with clade: Table of tree metadata by species (including clade number)
5. Chi squared summary/residuals: Table of clade-source statistics
6. Translation table: Conversion table between NCBI and GTDB reference genomes
7. Sankey translation: genome classifications between GTDB, GTDB+ and RefSeq
8. Walker strains paired data: Table of sequenced type-strains and correspondence to GTDB references
9. GTDB+ references pairwise ANI: A table which provide the pairwise ANI distances and alignment fraction between the reference genomes within GTDB+

#### 2 Distribution of ANI of isolates to all species within the GTDB schema

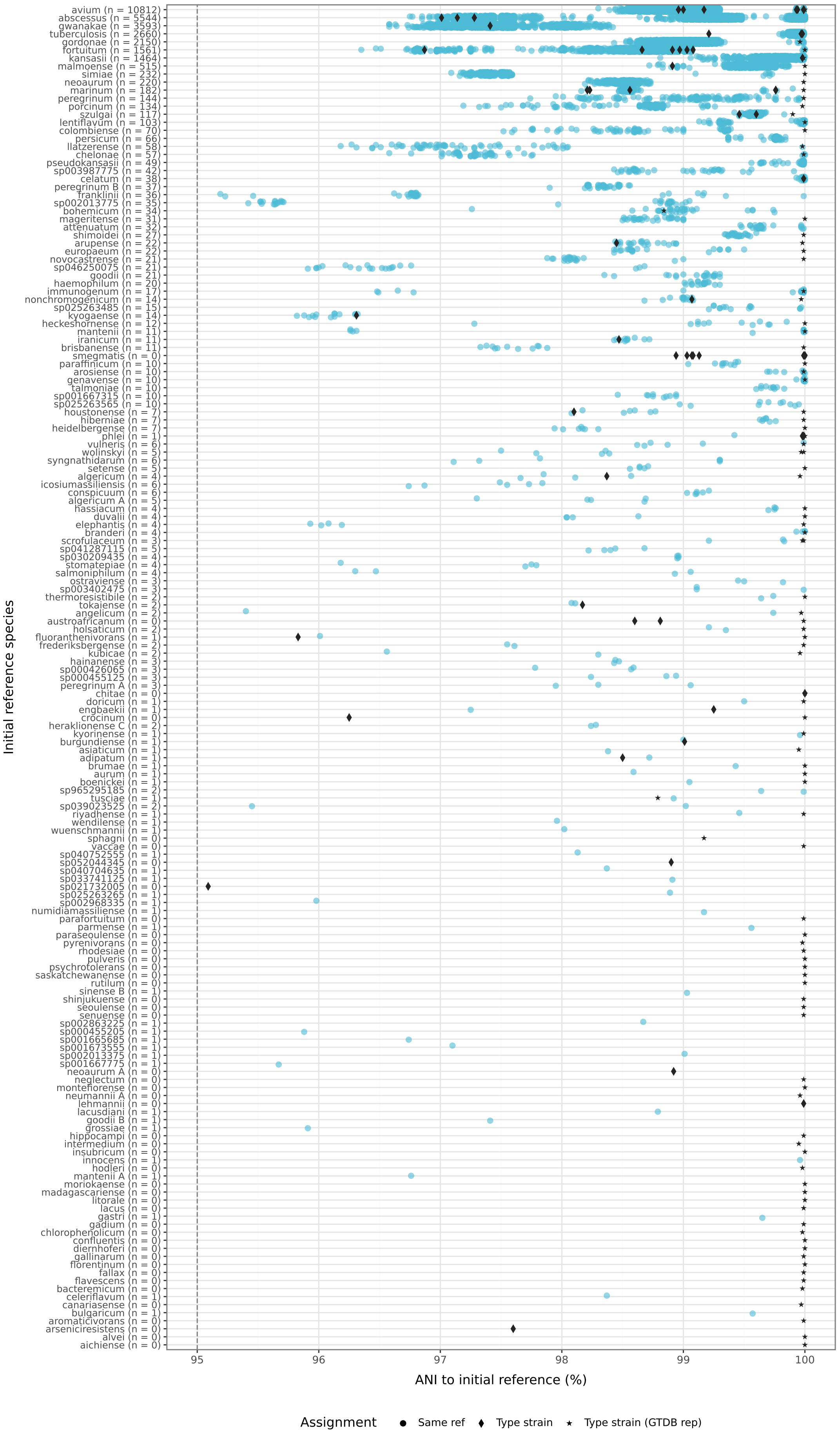

avium (n = 10812)
abscessus (n = 5544)
gwanakae (n = 3593)
tuberculosis (n = 2660)
gordonae (n = 2150)
fortuitum (n = 1561)
kansasii (n = 1464)
malmoense (n = 515)
simiae (n = 232)
neoaurum (n = 220)
marinum (n = 182)
peregrinum (n = 144)
porcinum (n = 134)
szulgai (n = 117)
lentiflavum (n = 103)
colombiense (n = 70)
persicum (n = 66)
llatzerense (n = 58)
chelonae (n = 57)
pseudokansasii (n = 49)
sp003987775 (n = 42)
celatum (n = 38)
peregrinum B (n = 37)
franklinii (n = 36)
sp002013775 (n = 35)
bohemicum (n = 34)
mageritense (n = 31)
attenuatum (n = 32)
shimoidei (n = 27)
arupense (n = 22)
europaeum (n = 22)
novocastrense (n = 21)
sp046250075 (n = 21)
goodii (n = 21)
haemophilum (n = 20)
immunogenum (n = 17)
nonchromogenicum (n = 14)
sp025263485 (n = 15)
kyogaense (n = 14)
heckeshornense (n = 12)
mantenii (n = 11)
iranicum (n = 11)
brisbanense (n = 11)
smegmatis (n = 0)
paraffinicum (n = 10)
arosiense (n = 10)
genavense (n = 10)
talmoniae (n = 10)
sp001667315 (n = 10)
sp025263565 (n = 10)
houstonense (n = 7)
hiberniae (n = 7)
heidelbergense (n = 7)
phlei (n = 1)
vulneris (n = 6)
wolinskyi (n = 5)
syngnathidarum (n = 6)
setense (n = 5)
algericum (n = 4)
icosiumassiliensis (n = 6)
conspicuum (n = 6)
algericum A (n = 5)
hassiacum (n = 4)
duvalii (n = 4)
elephantis (n = 4)
branderi (n = 4)
scrofulaceum (n = 3)
sp041287115 (n = 5)
sp030209435 (n = 4)
stomatepiae (n = 4)
salmoniphilum (n = 4)
ostraviense (n = 3)
sp003402475 (n = 3)
thermoresistibile (n = 2)
tokaiense (n = 2)
angelicum (n = 2)
austroafricanum (n = 0)
holsaticum (n = 2)
fluoranthenivorans (n = 1)
frederiksbergense (n = 2)
kubicae (n = 2)
hainanense (n = 3)
sp000426065 (n = 3)
sp000455125 (n = 3)
peregrinum A (n = 3)
chitae (n = 0)
doricum (n = 1)
engbaekii (n = 1)
crocinum (n = 0)
heraklionense C (n = 2)
kyorinense (n = 1)
burgundiense (n = 1)
asiaticum (n = 1)
adipatum (n = 1)
brumae (n = 1)
aurum (n = 1)
boenickei (n = 1)
sp965295185 (n = 2)
tusciae (n = 1)
sp039023525 (n = 2)
riyadhense (n = 1)
wendilense (n = 1)
wuenschmannii (n = 1)
sphagni (n = 0)
vaccae (n = 0)
sp040752555 (n = 1)
sp052044345 (n = 0)
sp040704635 (n = 1)
sp033741125 (n = 1)
sp021732005 (n = 0)
sp025263265 (n = 1)
sp002968335 (n = 1)
numidiamassiliense (n = 1)
parafortuitum (n = 0)
parmense (n = 1)
paraseoulense (n = 0)
pyrenivorans (n = 0)
rhodesiae (n = 0)
pulveris (n = 0)
psychrotolerans (n = 0)
saskatchewanense (n = 0)
rutilum (n = 0)
sinense B (n = 1)
shinjukuense (n = 0)
seoulense (n = 0)
senuense (n = 0)
sp002863225 (n = 1)
sp000455205 (n = 1)
sp001665685 (n = 1)
sp001673555 (n = 1)
sp002013375 (n = 1)
sp001667775 (n = 1)
neoaurum A (n = 0)
neglectum (n = 0)
montefiorense (n = 0)
neumannii A (n = 0)
lehmannii (n = 0)
lacusdiani (n = 1)
goodii B (n = 1)
grossiae (n = 1)
hippocampi (n = 0)
intermedium (n = 0)
insubricum (n = 0)
innocens (n = 1)
hodleri (n = 0)
mantenii A (n = 1)
moriokaense (n = 0)
madagascariense (n = 0)
litorale (n = 0)
lacus (n = 0)
gastri (n = 1)
gadium (n = 0)
chlorophenolicum (n = 0)
confluentis (n = 0)
diernhoferi (n = 0)
gallinarum (n = 0)
florentinum (n = 0)
fallax (n = 0)
flavescens (n = 0)
bacteremicum (n = 0)
celeriflavum (n = 1)
canariasense (n = 0)
bulgaricum (n = 1)
aromaticivorans (n = 0)
arseniciresistens (n = 0)
alvei (n = 0)
aichiense (n = 0)
Initial reference species
95
96
97
98
99
100
ANI to initial reference (%)
Assignment
Same ref
Type strain
Type strain (GTDB rep)

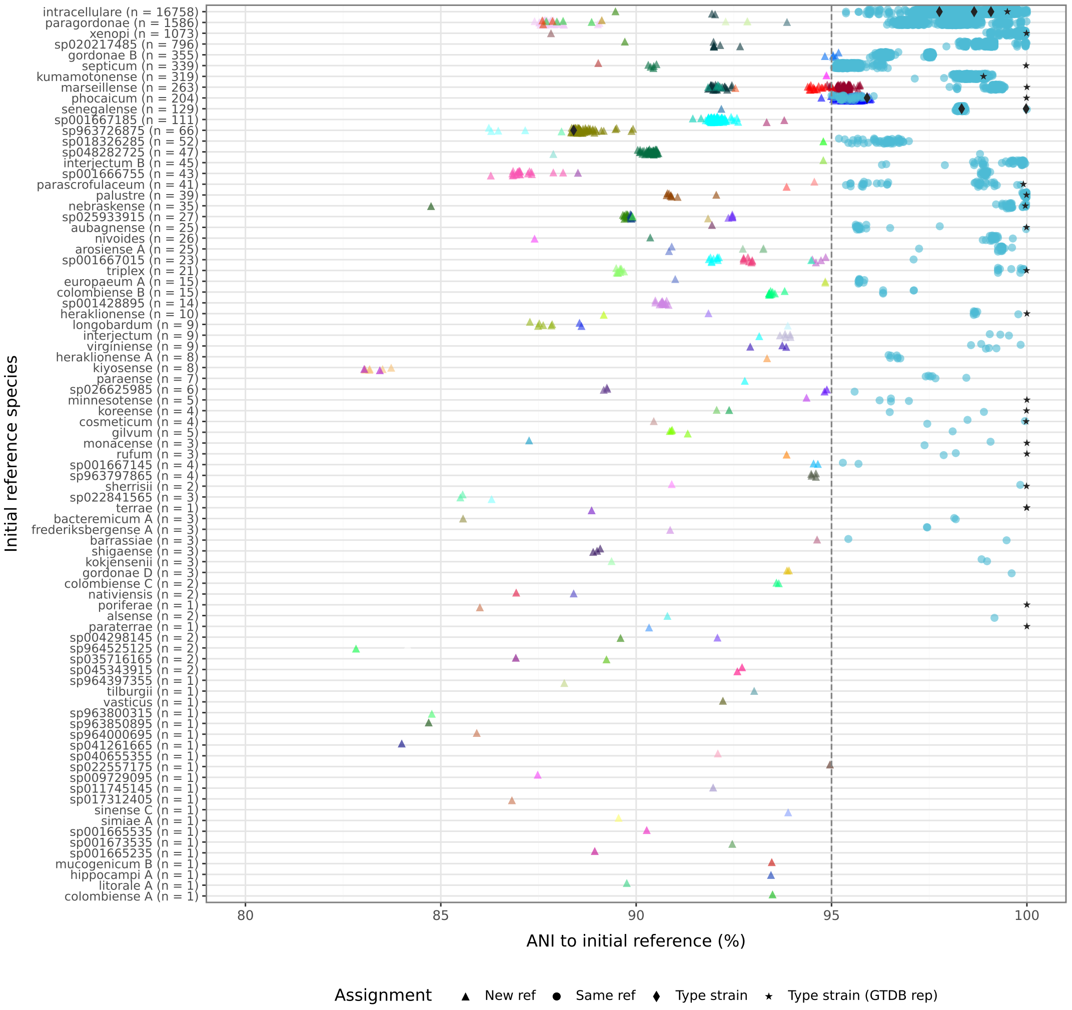


Figure S1: Plot of ANI distributions of isolates assigned to all GTDB reference species. Only GTDB species with at least one assigned isolate are shown. Panel A contains only species where all assigned isolates had ANI $\geq 95\%$. Panel B only species that had at least 1 isolate with an ANI $< 95$. Novel species-level genomic clusters are highlighted

### 3 Distribution of pairwise ANI between species reference genomes within GTDB+

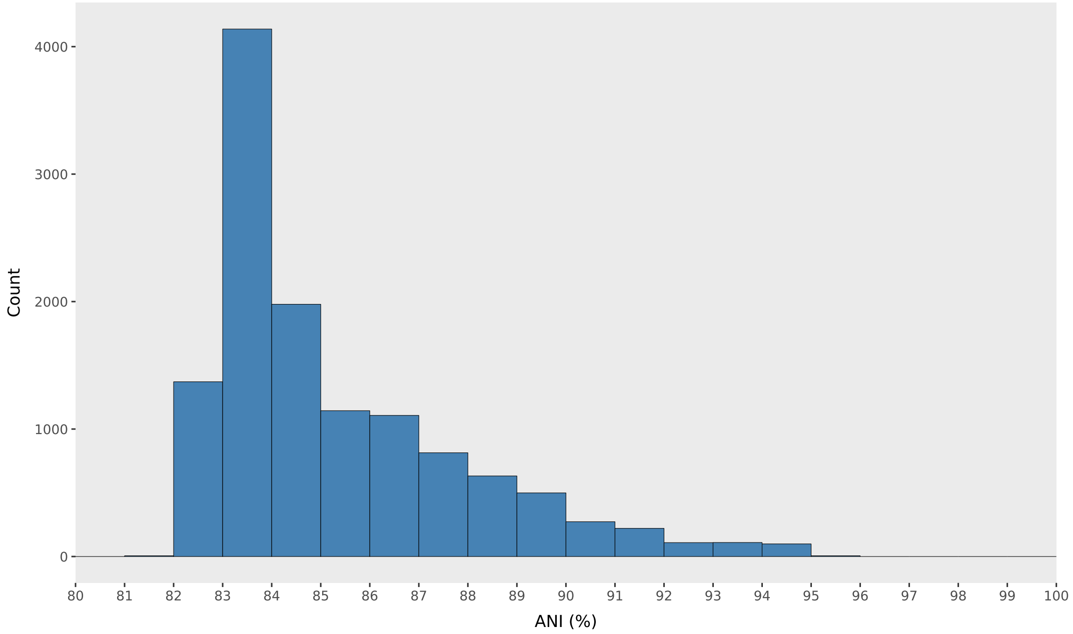


Figure S2: Distribution of pairwise average nucleotide identity (ANI) between the 960 GTDB+ reference genomes. Of the 460,320 possible pairwise comparisons, only seven (0.0015%) had ANI ≥95%. None of these seven comparisons involved the additional candidate species-level reference genomes identified in this study.

Table S1: Pairs of GTDB+ reference genomes with average nucleotide identity (ANI) ≥95%. Alignment fractions (AF) are reported for both the reference and query genomes. All seven pairs correspond to existing GTDB reference genomes; none involve the additional candidate species-level references identified in this study.

| Reference species | Query species | ANI (%) | AF ref. (%) | AF query (%) |
|---|---|---|---|---|
| *M. chelonae* | *M. gwanakae* | 95.18 | 85.90 | 85.37 |
| *M. sp002043095* | *M. hodleri_B* | 95.02 | 74.39 | 75.80 |
| *M. sp001667665* | *M. sp900240975* | 95.22 | 81.39 | 84.70 |
| *M. sp005789055* | *M. aquicola* | 95.32 | 80.93 | 53.15 |
| *M. sp020438785* | *M. sp020438865* | 95.05 | 70.47 | 57.09 |
| *M. chlorophenolicum* | *M. chubuense* | 95.53 | 69.32 | 86.05 |
| *M. sp963751315* | *M. sp052143115* | 95.00 | 61.53 | 60.07 |

### 4 Number of species by cluster size

Table S2: Number of all species-level clusters and additional candidate species-level clusters by minimum cluster size in the UKHSA collection.

| Minimum cluster size | All species | Novel species |
|---|---|---|
| 1 | 275 | 110 |
| 2 | 171 | 50 |
| 3 | 142 | 38 |
| 4 | 123 | 28 |
| 5 | 109 | 23 |
| 6 | 100 | 19 |
| 7 | 92 | 17 |
| 8 | 88 | 17 |
| 9 | 83 | 13 |
| 10 | 83 | 13 |

### 5 Phylogenetics

Table S3: Species used as outgroups for phylogenetic tree construction. Accession numbers refer to the genome assemblies used in the analysis.

| Species | Strain | Accession |
|---|---|---|
| *Corynebacterium diphtheriae* | NCTC 11397 | GCA_001457455.1 |
| *Gordonia bronchialis* | DSM 43247 | GCA_000024785.1 |
| *Nocardia farcinica* | NCTC 11134 | GCA_001182745.2 |
| *Rhodococcus erythropolis* | NBRC 15567 | GCA_001552595.1 |
| *Segniliparus rotundus* | DSM 44985 | GCA_000092825.1 |
| *Tsukamurella paurometabola* | DSM 20162 | GCA_000092225.1 |

Table S4: Fritz and Purvis D statistics for phylogenetic associations with ecological source and geographical region. Species without known source or geographical metadata were excluded from the corresponding analyses. A D statistic of 1 indicates a random phylogenetic distribution, whereas D values of 0 or below indicate phylogenetic clustering consistent with, or stronger than, the Brownian threshold model. P-values are reported against both random and Brownian null models. All calculations were performed using 10,000 permutations.

| Trait | D | p-value vs random | p-value vs Brownian | # with trait | # with data |
|---|---|---|---|---|---|
| UKHSA | 0.326 | 0 | 9e-04 | 275 | 931 |
| Human | 0.101 | 0 | 0.2029 | 399 | 931 |
| Environment | 0.339 | 0 | 0.0014 | 638 | 931 |
| Land environment | 0.437 | 0 | 0 | 453 | 931 |
| Domestic environment | 0.644 | 0 | 0 | 77 | 931 |
| Aquatic | 0.497 | 0 | 1e-04 | 161 | 931 |
| Freshwater | 0.581 | 0 | 0 | 93 | 931 |
| Saltwater | 0.576 | 0 | 0.0012 | 50 | 931 |
| East Asia & Pacific | 0.832 | 1e-04 | 0 | 222 | 908 |
| Europe & Central Asia | 0.722 | 0 | 0 | 560 | 908 |
| Latin America & Caribbean | 0.806 | 7e-04 | 0 | 65 | 908 |
| Middle East, North Africa, Afghanistan & Pakistan | 0.947 | 0.2818 | 0 | 27 | 908 |
| North America | 0.884 | 0.002 | 0 | 209 | 908 |
| South Asia | 0.611 | 0 | 0.0018 | 34 | 908 |
| Sub-Saharan Africa | 0.759 | 0 | 0 | 80 | 908 |

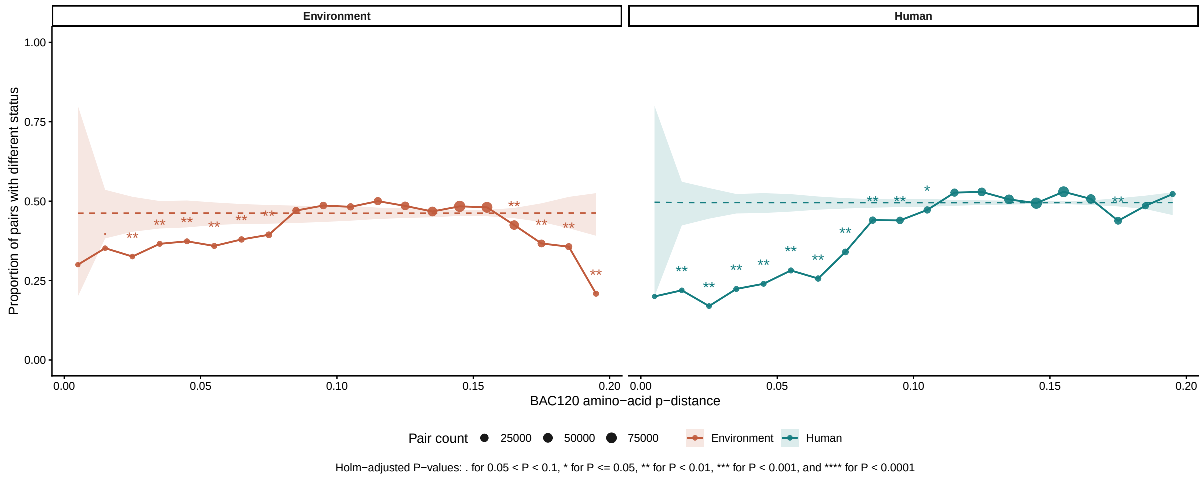


(a) Discordance of human and environmental sources of species across amino-acid p-distance.

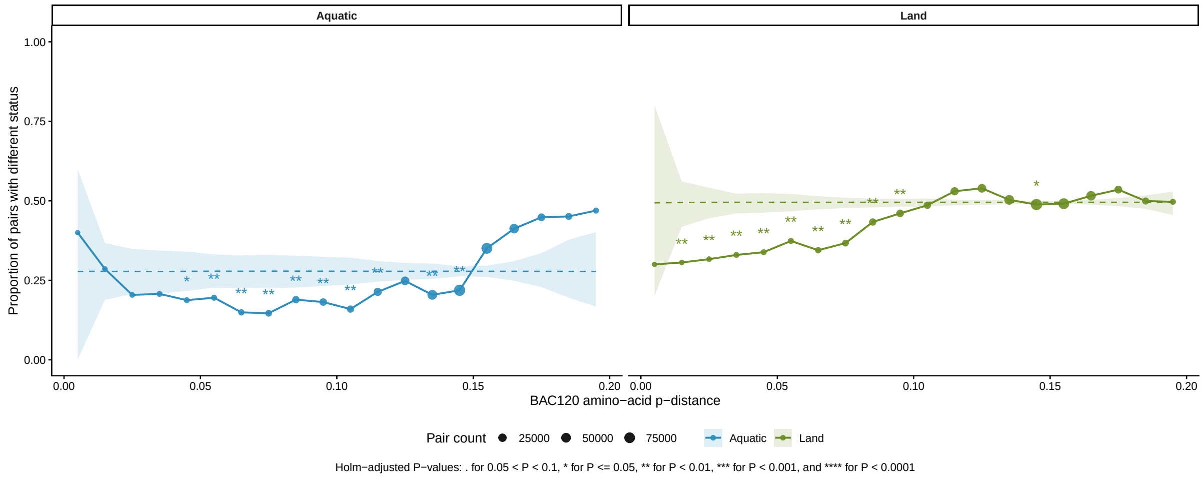


(b) Discordance of aquatic and land sources of species across amino-acid p-distance.

Figure S3: Discordance of species source classifications across amino-acid p-distance. (a) Human and environmental sources. (b) Aquatic and land sources. Pairwise p-distances among 887 high-quality reference genomes were grouped into 0.01-wide bins from 0 to 0.20. Points show the proportion of pairs with different source status; point size indicates the number of pairs within each bin. Dashed lines show the permutation-null mean, and shaded bands show the 2.5th–97.5th percentiles of the null distribution.

Table S5: Observed source concordance proportions by BAC120 amino-acid p-distance bin (1 = agreement, 0 = disagreement). The NULL row reports the pair-count-weighted permutation-null expectation. The number of pairwise comparisons per bin are shown in parentheses. Holm-corrected p-values are shown in square brackets.

| | Mean source concordance (n) | | | |
|---|---|---|---|---|
| AA p-distance bin | Human | Environment | Aquatic | Land |
| NULL | 0.505 [–] | 0.538 [–] | 0.722 [–] | 0.505 [–] |
| [0.00, 0.01) (n=10) | 0.800 [0.5391] | 0.700 [1.0000] | 0.600 [1.0000] | 0.700 [1.0000] |
| [0.01, 0.02) (n=196) | 0.781 [0.0020] | 0.648 [0.0600] | 0.714 [1.0000] | 0.694 [0.0020] |
| [0.02, 0.03) (n=436) | 0.830 [0.0020] | 0.674 [0.0032] | 0.796 [0.2088] | 0.683 [0.0020] |
| [0.03, 0.04) (n=1225) | 0.776 [0.0020] | 0.634 [0.0032] | 0.793 [0.2088] | 0.670 [0.0020] |
| [0.04, 0.05) (n=1167) | 0.760 [0.0020] | 0.626 [0.0032] | 0.812 [0.0198] | 0.662 [0.0020] |
| [0.05, 0.06) (n=1490) | 0.718 [0.0020] | 0.641 [0.0032] | 0.805 [0.0091] | 0.626 [0.0020] |
| [0.06, 0.07) (n=3041) | 0.744 [0.0020] | 0.621 [0.0032] | 0.851 [0.0020] | 0.655 [0.0020] |
| [0.07, 0.08) (n=5648) | 0.660 [0.0020] | 0.606 [0.0055] | 0.854 [0.0020] | 0.633 [0.0020] |
| [0.08, 0.09) (n=10948) | 0.560 [0.0020] | 0.530 [1.0000] | 0.811 [0.0091] | 0.567 [0.0020] |
| [0.09, 0.10) (n=13604) | 0.561 [0.0020] | 0.514 [1.0000] | 0.818 [0.0020] | 0.540 [0.0036] |
| [0.10, 0.11) (n=10285) | 0.528 [0.0100] | 0.518 [1.0000] | 0.840 [0.0020] | 0.514 [0.6093] |
| [0.11, 0.12) (n=21509) | 0.473 [1.0000] | 0.500 [1.0000] | 0.786 [0.0042] | 0.470 [1.0000] |
| [0.12, 0.13) (n=29502) | 0.471 [1.0000] | 0.515 [1.0000] | 0.752 [0.1360] | 0.461 [1.0000] |
| [0.13, 0.14) (n=51382) | 0.494 [1.0000] | 0.533 [1.0000] | 0.796 [0.0020] | 0.497 [1.0000] |
| [0.14, 0.15) (n=90591) | 0.506 [1.0000] | 0.516 [1.0000] | 0.781 [0.0020] | 0.512 [0.0154] |
| [0.15, 0.16) (n=79561) | 0.470 [1.0000] | 0.519 [1.0000] | 0.649 [1.0000] | 0.509 [0.3050] |
| [0.16, 0.17) (n=45624) | 0.493 [1.0000] | 0.575 [0.0020] | 0.587 [1.0000] | 0.484 [1.0000] |
| [0.17, 0.18) (n=16468) | 0.562 [0.0020] | 0.633 [0.0020] | 0.552 [1.0000] | 0.465 [1.0000] |
| [0.18, 0.19) (n=8466) | 0.515 [1.0000] | 0.643 [0.0020] | 0.549 [1.0000] | 0.501 [1.0000] |
| [0.19, 0.20) (n=1447) | 0.478 [1.0000] | 0.791 [0.0020] | 0.531 [1.0000] | 0.503 [1.0000] |

| Clade (number of species) | Human | Environment | Land | Aquatic |
|---|---|---|---|---|
| 18 (n=100) | 4.8 (3-7.7)<br><0.001 | 0.2 (0.13-0.3)<br><0.001 | 0.37 (0.23-0.58)<br><0.001 | 0.37 (0.18-0.78)<br>0.0053 |
| 11 (n=57) | 8.1 (3.9-17)<br><0.001 | 0.24 (0.13-0.41)<br><0.001 | 0.29 (0.15-0.54)<br><0.001 | 0.74 (0.35-1.6)<br>0.59 |
| 6 (n=50) | 6.7 (3.2-14)<br><0.001 | 0.36 (0.2-0.64)<br><0.001 | 0.43 (0.23-0.8)<br>0.0082 | 0.043 (0.0027-0.71)<br><0.001 |
| 30 (n=42) | 7.3 (3.2-17)<br><0.001 | 0.43 (0.23-0.8)<br>0.0096 | 0.57 (0.3-1.1)<br>0.084 | 1.7 (0.83-3.4)<br>0.15 |
| 15 (n=37) | 3.8 (1.8-8)<br><0.001 | 0.18 (0.085-0.36)<br><0.001 | 0.32 (0.15-0.69)<br>0.0023 | 0.26 (0.061-1.1)<br>0.047 |
| 4 (n=6) | 16 (0.91-290)<br>0.0059 | 0.088 (0.01-0.76)<br>0.012 | 0.21 (0.024-1.8)<br>0.22 | 0.38 (0.021-6.9)<br>0.6 |
| 32 (n=16) | 3 (1-8.8)<br>0.041 | 0.74 (0.27-2.1)<br>0.59 | 0.62 (0.22-1.7)<br>0.45 | 1.1 (0.3-3.8)<br>1 |
| 17 (n=37) | 2 (1-4)<br>0.041 | 0.46 (0.24-0.88)<br>0.028 | 0.7 (0.36-1.4)<br>0.32 | 0.72 (0.27-1.9)<br>0.66 |
| 1 (n=15) | 2.7 (0.93-8.1)<br>0.068 | 6.4 (0.84-49)<br>0.047 | 1.2 (0.43-3.3)<br>0.8 | 7.3 (2.6-21)<br><0.001 |
| 31 (n=29) | 1.7 (0.8-3.5)<br>0.18 | 1.4 (0.6-3.4)<br>0.54 | 1.1 (0.54-2.4)<br>0.85 | 2.5 (1.2-5.6)<br>0.024 |
| 8 (n=18) | 1.7 (0.66-4.4)<br>0.34 | 0.7 (0.27-1.8)<br>0.45 | 1.3 (0.51-3.4)<br>0.64 | 0.27 (0.035-2)<br>0.22 |
| 29 (n=16) | 1.7 (0.65-4.7)<br>0.31 | 0.74 (0.27-2.1)<br>0.59 | 1 (0.39-2.8)<br>1 | 0.66 (0.15-2.9)<br>0.75 |
| 26 (n=5) | 0.9 (0.15-5.4)<br>1 | 4.5 (0.25-83)<br>0.33 | 1.6 (0.26-9.4)<br>0.68 | 1.2 (0.13-10)<br>1 |
| 16 (n=9) | 0.67 (0.17-2.7)<br>0.74 | 0.9 (0.22-3.6)<br>1 | 2.1 (0.52-8.5)<br>0.33 | 0.25 (0.015-4.4)<br>0.37 |
| 27 (n=7) | 0.54 (0.1-2.8)<br>0.71 | 1.1 (0.22-5.8)<br>1 | 1.4 (0.31-6.3)<br>0.72 | 3.5 (0.78-16)<br>0.11 |
| 20 (n=31) | 0.63 (0.29-1.4)<br>0.27 | 1.9 (0.77-4.7)<br>0.17 | 0.98 (0.48-2)<br>1 | 1.1 (0.45-2.8)<br>0.81 |
| 28 (n=55) | 0.7 (0.39-1.2)<br>0.26 | 1.9 (0.94-3.6)<br>0.073 | 0.93 (0.54-1.6)<br>0.89 | 2 (1.1-3.7)<br>0.029 |
| 24 (n=79) | 0.64 (0.39-1)<br>0.074 | 1.2 (0.7-2)<br>0.61 | 1.8 (1.1-2.9)<br>0.018 | 0.23 (0.082-0.63)<br>0.001 |
| 33 (n=7) | 0.095 (0.0054-1.7)<br>0.023 | 6.3 (0.36-110)<br>0.11 | 2.6 (0.51-14)<br>0.28 | 1.9 (0.36-9.7)<br>0.36 |
| 14 (n=7) | 0.095 (0.0054-1.7)<br>0.023 | 6.3 (0.36-110)<br>0.11 | 1.4 (0.31-6.3)<br>0.72 | 1.9 (0.36-9.7)<br>0.36 |
| 7 (n=9) | 0.073 (0.0042-1.3)<br>0.012 | 8.2 (0.47-140)<br>0.064 | 8.5 (1.1-68)<br>0.019 | 0.25 (0.015-4.4)<br>0.37 |
| 34 (n=10) | 0.066 (0.0038-1.1)<br>0.0065 | 9.1 (0.53-160)<br>0.036 | 21 (1.2-370)<br><0.001 | 0.23 (0.013-3.9)<br>0.22 |
| 12 (n=17) | 0.082 (0.011-0.62)<br>0.0019 | 7.3 (0.97-56)<br>0.03 | 5 (1.4-18)<br>0.006 | 0.99 (0.28-3.5)<br>1 |
| 35 (n=30) | 0.2 (0.069-0.57)<br>0.001 | 4.2 (1.3-14)<br>0.0089 | 2.5 (1.1-5.5)<br>0.025 | 0.71 (0.24-2.1)<br>0.63 |
| 2 (n=19) | 0.034 (0.0021-0.57)<br><0.001 | 18 (1.1-290)<br>0.0016 | 9.2 (2.1-40)<br><0.001 | 0.54 (0.12-2.4)<br>0.55 |
| 23 (n=36) | 0.12 (0.035-0.38)<br><0.001 | 5.1 (1.6-17)<br>0.0015 | 5.5 (2.3-13)<br><0.001 | 0.13 (0.017-0.93)<br>0.013 |
| 21 (n=35) | 0.077 (0.018-0.32)<br><0.001 | 7.8 (1.8-33)<br><0.001 | 1.6 (0.8-3.2)<br>0.23 | 1.6 (0.75-3.6)<br>0.26 |
| 9 (n=68) | 0.095 (0.038-0.24)<br><0.001 | 11 (3.3-34)<br><0.001 | 15 (6-38)<br><0.001 | 0.13 (0.031-0.54)<br><0.001 |
| 22 (n=61) | 0.0097 (6e-04-0.16)<br><0.001 | 61 (3.7-980)<br><0.001 | 0.083 (0.033-0.21)<br><0.001 | 22 (12-40)<br><0.001 |

Corrected odds ratio (95% CI) and Fisher's exact test p-value

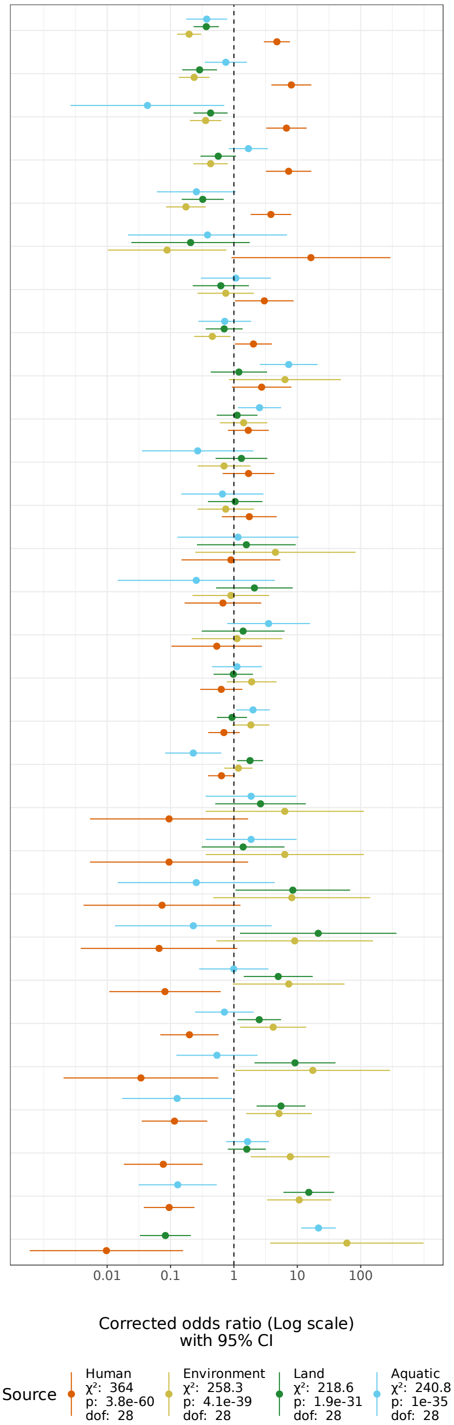


Figure S4: Odds ratio for source types within clades.

a) Table of corrected odds ratio with 95% confidence intervals and fisher exact p-values. Squares are coloured based on standardised Pearson residuals.

b) Forest plot of corrected odds ratio with 95% confidence intervals.

The $\chi^2$-statistic, p-value and degrees of freedom are provided in the legend. Clades have been ordered by the residuals for "Human". Note that a given species may be found from multiple sources. Aquatic (oceans/rivers/fish) and Land (soil/plants/animals) are subsets of Environment.

### 6 Rarefaction curve of observed species-level clusters

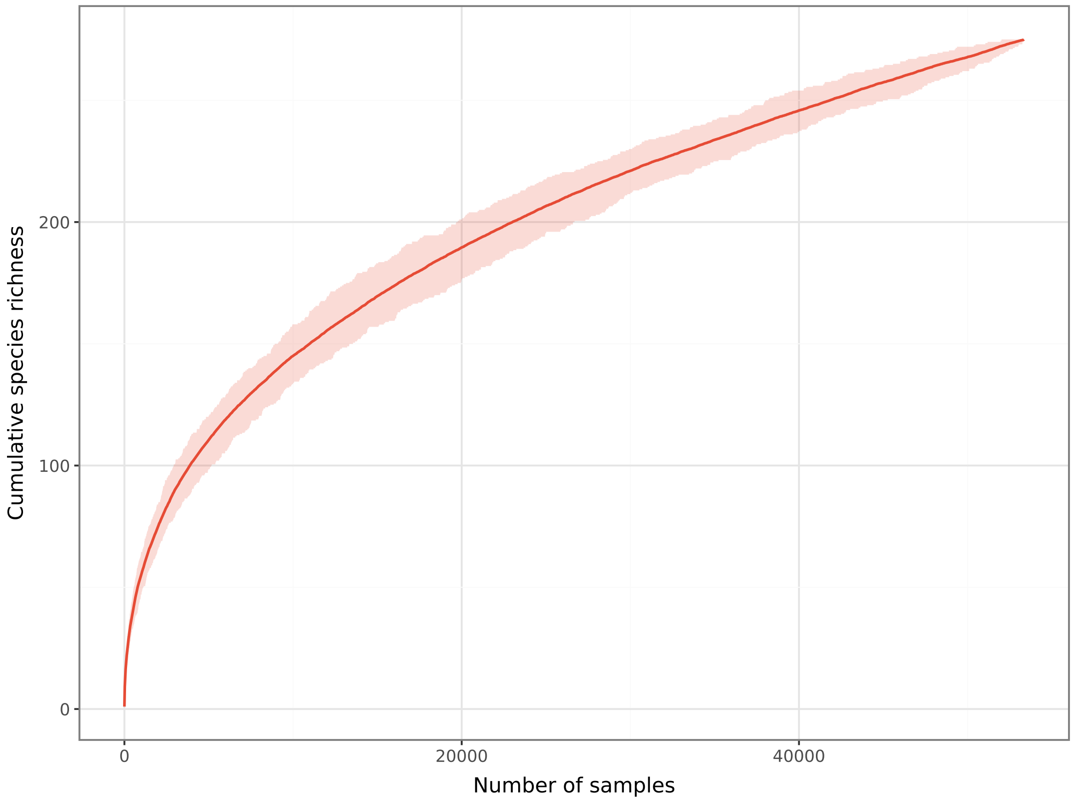


Figure S5: A rarefaction curve showing the accumulation of *Mycobacterium* species as a function of sample size. The solid red line denotes the mean number of unique GTDB+ species observed at each sampling depth, with the 95% quantile interval shown in shaded red.
